# Missense mutations in the calcium-activated chloride channel TMEM16A promote tumor growth by activating oncogenic signaling in Human Cancer

**DOI:** 10.1101/2023.06.21.545912

**Authors:** Silvia Cruz-Rangel, Jose Juan De Jesus-Perez, Avani Gopalkrishnan, Roberto Gomez-Casal, Jonathan Pacheco, Maya R Brown, Abdulkader Yassin-Kassab, Gerald RV Hammond, Carol A Bertrand, Jorge Arreola, Kirill Kiselyov, Duvvuri Umamaheswar

## Abstract

The calcium-activated chloride channel TMEM16A is overexpressed in several tumors. This condition is associated with a poor survival prognosis but highlights TMEM16A’s potential as a biomarker and target for anti-cancer therapies. Numerous somatic mutations of TMEM16A have been reported; however, their potential and molecular mechanism of oncogenesis are unknown. Here, we investigate the function and oncogenicity of nine-point mutations found in human cancerous tissues (R451P, R455Q, M546I, R557W, F604L, D902N, K913E, D914H, and Q917K). These mutations are located on the extracellular side and near the third Ca^2+^-binding site, near a PtdIns(4,5)P2 site in the human TMEM16A channel. Our findings reveal that these mutations affected gating, Ca^2+^ sensitivity, phosphorylation of essential signaling proteins, cell proliferation, and tumor growth. Notably, R451P and D902N exhibit low Ca^2+^ sensitivity, yet their overexpression promotes phosphorylation of EGFR and AKT, as well as *in vivo* tumorigenesis, without Ca^2+-^enhancing stimuli. Conversely, the charged-neutralizing mutation R451Q and the conservative mutation D902E restored Ca^2+^ sensitivity and altered cell proliferation and tumor growth as wild-type did. Thus, we conclude that the oncogenic phenotype of TMEM16A missense mutations is independent of chloride flux but involves the differential activation of cell signaling components associated with cell proliferation.

## INTRODUCTION

The calcium-activated chloride (Cl^-^) channel TMEM16A is an extensively expressed protein that supports a wide range of vital biological processes. For example, it serves as a critical player in cell proliferation and migration, fluid secretion in exocrine glands, regulation of neuronal and cardiac excitability, nociception, maintenance of smooth muscle tone, facilitation of gut motility, modulation of apoptosis, and regulation of cell volume (For recent reviews see: (1-5)). Numerous structure-function studies combined with Cryo-EM assays and molecular dynamics analysis have started to unveil the intricate molecular mechanisms underlying the activation and regulation of TMEM16A by both intra-and extracellular stimuli. The activation of TMEM16A requires the coincidence of two key factors: intracellular calcium (Ca^2+^) and membrane depolarization. Complete activation of TMEM16A occurs when the concentration of intracellular calcium ([Ca^2+^]_i_) surpasses 1.5 µM (6-11). Channe’s activity is modulated by intra-and extracellular protons (12-15) and the permeating anion, which seems to couple the gating and permeation processes (16-19). Furthermore, the TMEM16A functionality is strongly influenced by its interaction with various membrane lipids, including cholesterol, fatty acids, and phosphatidylinositol 4,5-bisphosphate (PtdIns(4,5)P2) (20-28). TMEM16A is overexpressed in a diverse range of cancer tissues, including but not limited to oesophagal squamous cell carcinoma, head and neck squamous cell carcinoma, breast cancer, gastrointestinal stromal tumors, prostate cancer, colorectal cancer, hepatocellular carcinoma, glioma, and lung cancer (reviewed in (29-31)). The upregulation of TMEM16A protein expression has been found to negatively correlate with survival of patients diagnosed with malignant tumors (32, 33). Multiple hypotheses have been put forward to explain the potential role of TMEM16A in cancer development. These hypotheses encompass a broad spectrum of molecular targets closely associated with the overexpression of TMEM16A, which influence vital processes such as cell proliferation, migration, and invasiveness. Notably, EGFR and Her2, the Mitogen-activated protein kinase (MAPK), Ca^2+^/Calmodulin-dependent protein kinase II (CAMKII), and the Nuclear factor κB (NFκB) signaling pathways involving tyrosine kinase receptors have been implicated as potential targets of the TMEM16A overexpression in cancer development (1, 29, 31). On the other hand, the role of the Cl^-^ conductance, ostensibly the core function of TMEM16A, in tumor growth and cell proliferation remains undefined. Presumably, cancer cells overexpressing TMEM16A generate changes in the membrane potential associated with an enhanced Cl^-^ flux, which would alter fundamental signaling pathways controlling the cell proliferation rate. However, existing data particularly in various cancer cells have yielded contradictory results. For example, TMEM16A-dependent cell proliferation, migration and/or invasiveness across multiple cancer types are inhibited by channel function blockers such as T16Ainh-A01 or CaCCinh-A01 with similar efficacy as gene silencing strategies (34-40). Furthermore, as has been described by our group and others (35, 41, 42), the overexpression of defective Cl^-^ conductance TMEM16A mutants fails to replicate the oncogenic impact of the wild-type (WT) channel, supporting the significance of Cl^-^ conductance in tumorigenesis. However, contrary to these findings, the pharmacological inhibition of TMEM16A currents in pancreatic and colorectal cancer cell lines fails to reproduce the effects observed from silencing TMEM16A expression using siRNA on cell proliferation and viability (43, 44). Furthermore, Because TMEM16A interacts with proteins involved in cell growth and proliferation and EGFR (45), not obviously regulated by the intracellular Cl^-^ concentration ([Cl^-^]_i_) or the membrane potential, the oncogenic effect of TMEM16A could be mediated by turning on signaling proteins rather than by increasing the total Cl^-^ conductance.

Parsing data from The Cancer Genome Atlas (TCGA) shows the presence of 390 missense mutations in TMEM16A, whose effects on the carcinogenic process are unknown. Nine of these mutants (R451P, R455Q, M546I, R557W, F604L, D902N, K913E, D914H, and Q917K) are located on the extracellular side of the membrane, within the region of the channel associated with Cl^-^ selectivity and in a cluster of amino acids found in the vicinity of the recently characterized third Ca^2+^-binding site that allosterically regulates the canonical site responsible for channel activation (46). These mutations have been identified in patient tumor samples obtained from a wide range of cancer types, including Head and Neck Squamous Cell Carcinoma, Cutaneous Melanoma, Stomach Adenocarcinoma, Thymoma, Uterine Endometrioid Carcinoma, Prostate Adenocarcinoma, Glioblastoma Multiforme, and Lung Adenocarcinoma (as presented in Table 1). Thus, in this work, we used electrophysiology and *in vivo* and *in vitro* models to determine the impact of the nine missense mutations mentioned above in their Cl^-^ conductance and Ca^2+^ dependence. Additionally, we determine their effects on cell proliferation, tumor growth and oncogenic signaling without stimulating the cytosolic Ca^2+^ increase needed to activate the TMEM16A-dependent Cl^-^ conductance.

**Table 1.**
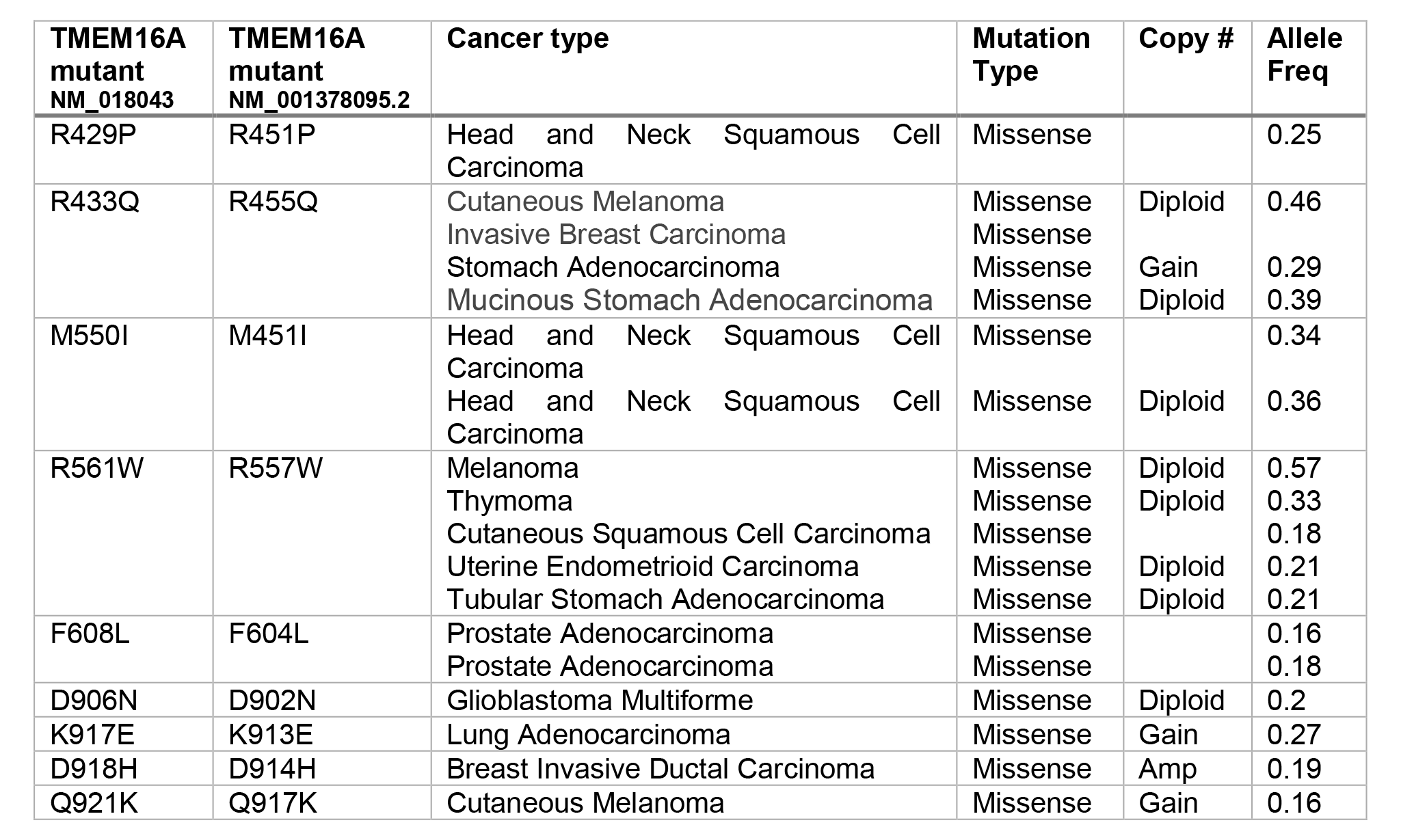
TMEM16A mutations described in cancer patient tumor samples evaluated in this work. Mutations are reported in the cBioPortal platform using hTMEM16A NM_018043 as a template.

Surprisingly, R451P, R557W, and D902N mutants have decreased their Ca^2+^ sensitivity but promoted cancer cell growth *in vitro* and *in vivo* better than the WT protein. Yet, the K913E somatic mutation, which exhibits a Ca^2+^-independent function, significantly reduced cell proliferation and tumor growth. Moreover, these mutations selectively stimulate distinct signaling pathways associated with cellular growth and proliferation. Specifically, the D902N mutant exhibits EGFR heightened activation, while the R451P mutant enhances AKT protein activity. Thus, low Ca^2+^ sensitive missense mutations can induce cell proliferation and tumor growth, most likely independently of Cl^-^ conductance. In contrast, the K913E mutant, characterized by a Ca^2+^-independent Vm activation, causes a notable decrease in proliferative capacity and tumor growth. Our data highlight the complexity of the mechanism underlying TMEM16A-dependent tumorigenesis, providing insights into the pivotal role of TMEM16A as a central molecule within oncogenic signaling pathways, functioning independently of its chloride conductance function.

## RESULTS

### The somatic mutations in the human TMEM16A channel are associated with cancer

Figure 1A shows the incidence of nonsense mutations in the entire human TMEM16A (hTMEM16A) or human ANO 1 coding sequence obtained from the cBioPortal platform (46, 47). Using a curated set of 191 non-redundant studies of DNA genomic sequencing from cancer patient samples (Supplemental document 1), 460 human mutations were identified (Figure 1A) scattered throughout the protein and characterized by their relative abundances. The mutations are divided into those that generate truncated proteins (38 mutants, black dots), mutations that occur at the limit of an exon and an intron (alternative splicing sites, 32 mutants, pink dots), and the most abundant missense mutations (390 mutants, green dots). To date, no data describing their effects on TMEM16A activation and function or data to rule in or rule out any of these mutations on carcinogenesis are available.

**FIGURE 1.**
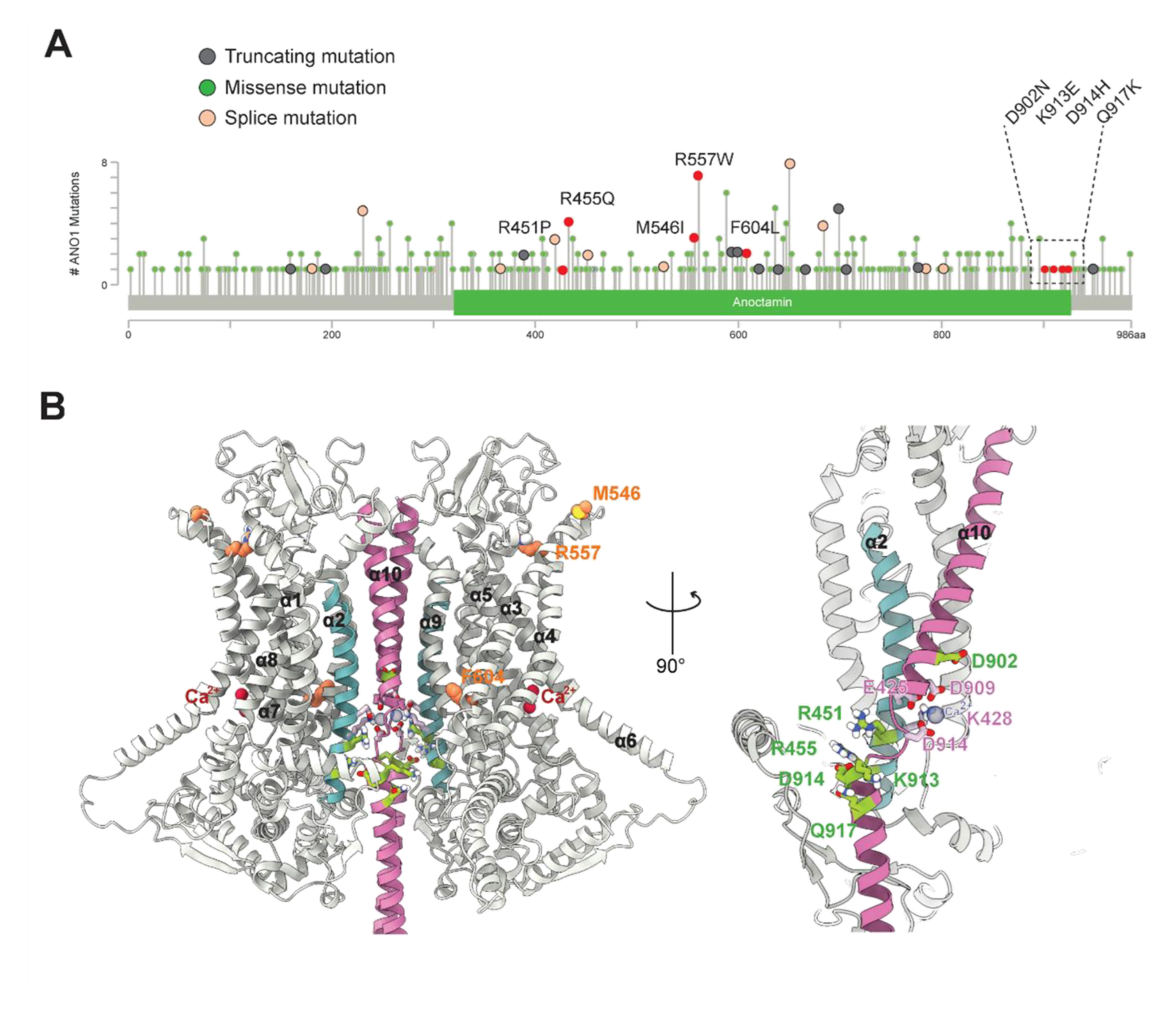
Somatic mutations in the human TMEM16A channel reported in the cBioPortal Cancer Genomics platform. **(A)** The lollipop spheres indicate the location and frequency of the TMEM16A mutations. Mutations analyzed in this work are shown in red in the TMEM16A lollipop plot. **(B)** Ribbon representation of hTMEM16A structure with the extracellular and intracellular sides at the top and bottom, respectively. The homology model was obtained from the AlphaFold Protein Structure Database. The left panel shows the dimer structure with the putative location of residues M546I, R557W, and F604L in orange. Red spheres in the left panel represent Ca^2+^ ions in the canonic intracellular Ca^2+^-binding site. The right panel highlight the helices α2, α6, and α10 from one subunit obtained by rotating the left structure 90˚clockwise around the two-fold axis to show the location of residues R451, R455, D902, K913, D914 and Q917 in green. Residues forming the third Ca^2+^ binding site between α2 and α10 are in pink, and the purple spheres represent a Ca^2+^ ion in this site.

We analyzed the effects of nine mutations on TMEM16A Ca^2+^ sensitivity, phosphorylation of proteins participating in cell proliferation, cell proliferation, and tumor growth. To map the putative location of the mutated residues, we used an Alpha Fold model structure of hTMEM16A (Figure 1B). The left panel shows the homodimeric hTMEM16A structure, with each subunit separated by their α10 helices at the center, forming a pseudo-symmetry axis. hTMEM16A mutation M546I is located at the extracellular end of TMD 3, R557W is at TMD 7, and F604L is at the N-terminus of the transmembrane domain 3. The right panel of Figure 1B shows the α2, α6, and α10 helices of one subunit turned 90^0^ clockwise to show that the cluster of amino acids R451P, R455Q, D902N, K913E, D914H and Q917H are located closer to the third Ca^2+^- binding site (Le and Yang, 2020). These intracellular residues are at the interfacial helixes connecting TMD2 and TMD10, close to a pocket identified as both an intracellular Ca^2+^-binding site and PtdIns(4,5)P2 binding domain (46). R451 is adjacent to K450 (K418 in mTMEM16A), while D902 is upstream of D905 (D879 in mTMEM16A); both residues coordinate Ca^2+^ binding in the third intracellular Ca^2+^ binding site (46). In addition, the sidechains of R451 form stable salt–bridge with residue K567 to coordinate PtdIns(4,5)P2 binding (Le et al., 2019). D902N is located at the intracellular end of TMD10, facing TMD9 of the same monomer and the same TMD9 of the adjacent subunit, probably forming a dynamic salt bridge with the K795 in its own TMD9 (PDB: 5OYB, 5OYG, 7ZK3).

In addition to their molecular location, their expression pattern suggests they could impact the development of various cancers. hTMEM16A M546I mutation is expressed in Head and Neck Squamous Cell Carcinoma (48-58), R557W/Q in Melanoma, Thymoma, Cutaneous Squamous Cell Carcinoma, Uterine Endometrium Carcinoma, and Tubular Stomach Adenocarcinoma samples (49-60); F604L was described in Prostate Adenocarcinoma (61), R455Q is a high-incidence mutation; its expression profile in human tissue includes Cutaneous Melanoma, Invasive Breast Carcinoma, Stomach Adenocarcinoma, and Mucinous Stomach Adenocarcinoma (49-53, 55-58, 61), R451P, K913E, D914H and Q917K were identified with a minor incidence in Head and Neck Squamous cell carcinoma, Lung Adenocarcinoma, Breast Invasive Ductal Carcinoma and Cutaneous Melanoma, respectively (48-58), and D902N in Glioblastoma multiform (46, 49-53, 55-58). A summary of mutants, including their genetic status in the tumors, is found in Table 1.

### Cancer-associated mutations alter chloride currents and calcium sensitivity of human TMEM16A

We reasoned that the Ca^2+^ sensitivity and the Cl^-^ conductance of TMEM16A are essential for its role in oncogenesis. If so, we will find a correlation between altered Ca^2+^ sensitivity and oncogenesis. To comprehensively examine this idea, we employed the hTMEM16A clone NM_001378095.2 to generate constructs containing cancer-associated point mutations. These constructs were transiently transfected into HEK293 cells, enabling us to conduct thorough functional analyses using a patch clamp.

A confocal localization analysis was conducted on HEK293 cells expressing GFP-tagged TMEM16A channels to verify the appropriate localization of the mutant variants. Cells transfected with the GFP-fused TMEM16A channels exhibited a fluorescent signal consistent with plasma membrane localization (Figure 2A). This observation indicates that none of the nine-point mutations had their localization altered because they properly transit and reach the cell membrane.

**FIGURE 2.**
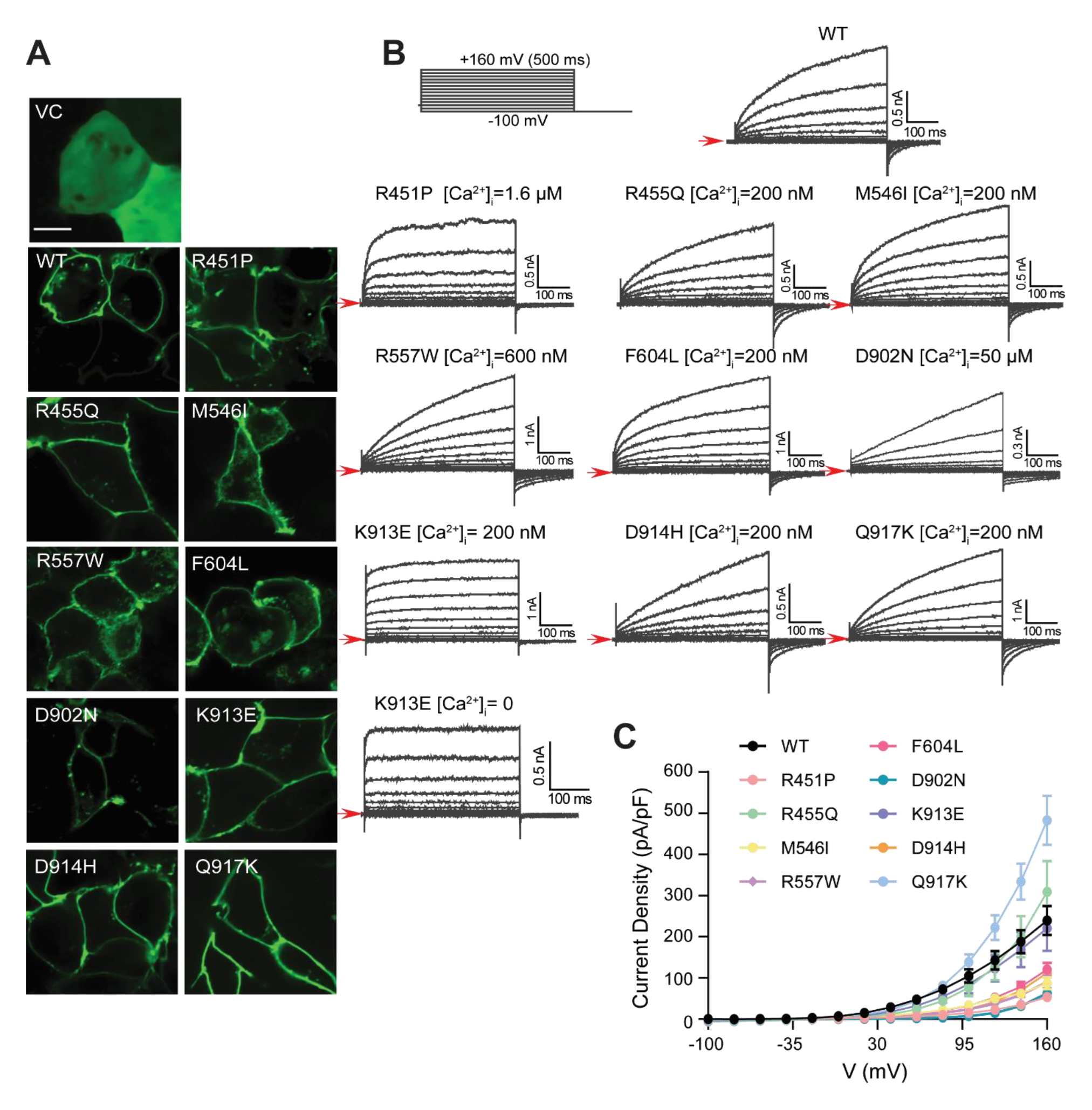
Cancer-associated mutations alter TMEM16A channel chloride currents. **(A)** Confocal analysis of TMEM16A channels expression. Representative confocal images through the equatorial plane of HEK293 cells transiently transfected with empty vector (pCMV6-EGFP), TMEM16A WT or the mutants (green). Images are maximum projections of a z stack of x optical slices. Scale bar 8 μm. **(B)** Representative macroscopic Cl^-^ currents produced by activation of TMEM16A WT and mutants. Whole cell currents were recorded in the presence of 0.2 µM (WT, R455Q, M546I, F604L, D918H and Q921K), 1.4 µM (R451P), 0.6 µM (R557W), 50 µM (D902N), and 0 nM (K913E) intracellular free Ca^2+^ concentration. The stimulation protocol is shown in the upper panel. The Red arrows indicate a 0-nA baseline. An amplification of the current around the reversal potential is shown (**C)** Mean steady-state whole-cell Cl^-^ current density-voltage relationships for WT and mutant channels (n= 6-13).

Next, we used whole-cell patch clamp recordings to evaluate the channel activity. Representative Cl^-^ currents (I_Cl_) generated by the WT and the nine mutants are shown in Figure 2B. The [Ca^2+^]_i_ necessary to induce activation varies among these proteins. Mutants R455Q, M546I, F604L, D914H, and Q917K caused outwardly rectifying I_Cl_ in the presence of 0.2 µM Ca^2+^, the same [Ca^2+^]_i_ used to activate the WT channel. The R451P mutant channel notably exhibits activation at significantly higher [Ca^2+^]_i_ levels (1.6 µM). This suggests a reduced Ca^2+^ sensitivity compared to the WT channel. On the other hand, the R567W mutant required a slight increase in [Ca^2+^]_i_ (0.6 µM) to induce its activation. Remarkably, the D902N mutant necessitates a considerably higher [Ca^2+^]_i_ (50 µM) to exhibit activation. Interestingly, the K913E mutant was active at 0.2 µM Ca^2+,^ but turned to a voltage-activated channel in cells dialyzed with a free-intracellular Ca^2+^ solution. The voltage-induced currents displayed strong o rectification in the absence or presence of 0.2 µM Ca^2+^. Therefore this mutant is Ca^2+^-independent or requires incredibly low [Ca^2+^]_i_ to be active. The range of current amplitudes of the R451P and D902N mutants were consistently small, even when high [Ca^2+^]_i_ were used (Figure 2B, C).

The previous observations suggested that some of these variants displayed altered Ca^2+^ sensitivity. We constructed concentration-response curves to Ca^2+^ in excised patches to examine this idea. Representative traces are shown in Figure 3A. The [Ca^2+^]_i_ needed to produce half of the maximum activation (EC_50_) of TMEM16A WT was 0.53 ± 0.13, and 1.42 ± 0.37 μM, at +80 and −80 mV, respectively. Consistent with our hypothesis, the EC_50_ of R557W at +80 mV was 1.33 ± 0.16 µM and 3.3 ± 0.16 µM at −80 mV (Figure 3B and 3C), confirming the reduced Ca^2+^ sensitivity of this mutant. R451P and D902N changed their apparent Ca^2+^ sensitivity even more than R557W. Dose-response curves in Fig 3B show that the EC_50_ for Ca^2+^ of the R451P channel is 6.9 ± 0.5 at +80 and 12.7 ± 2 μM at −80 mV, almost one order of magnitude less sensitive than WT TMEM16A. An even more drastic reduction of Ca^2+^ sensitivity was seen with the D902N mutant, with an EC_50_ of 30.2 ± 4.5 µM at +80 mV. Because of the small I_Cl_ amplitude of D902N and strong rectification at high [Ca^2+^]_i_ (Figure 2B, C), it was not possible to determine its EC_50_. Based on low Ca^2+^ sensitivity and outward rectification, we concluded that the D902N mutation could not allow the flux of Cl^-^ in any direction. Even with the Ca^2+^ sensitivity shift, these channels maintain the outward rectification. These findings highlight a crucial and hitherto undisclosed function of residues R451 and D902, influencing intracellular Ca^2+^ sensitivity of the TMEM16A channel.

**FIGURE 3.**
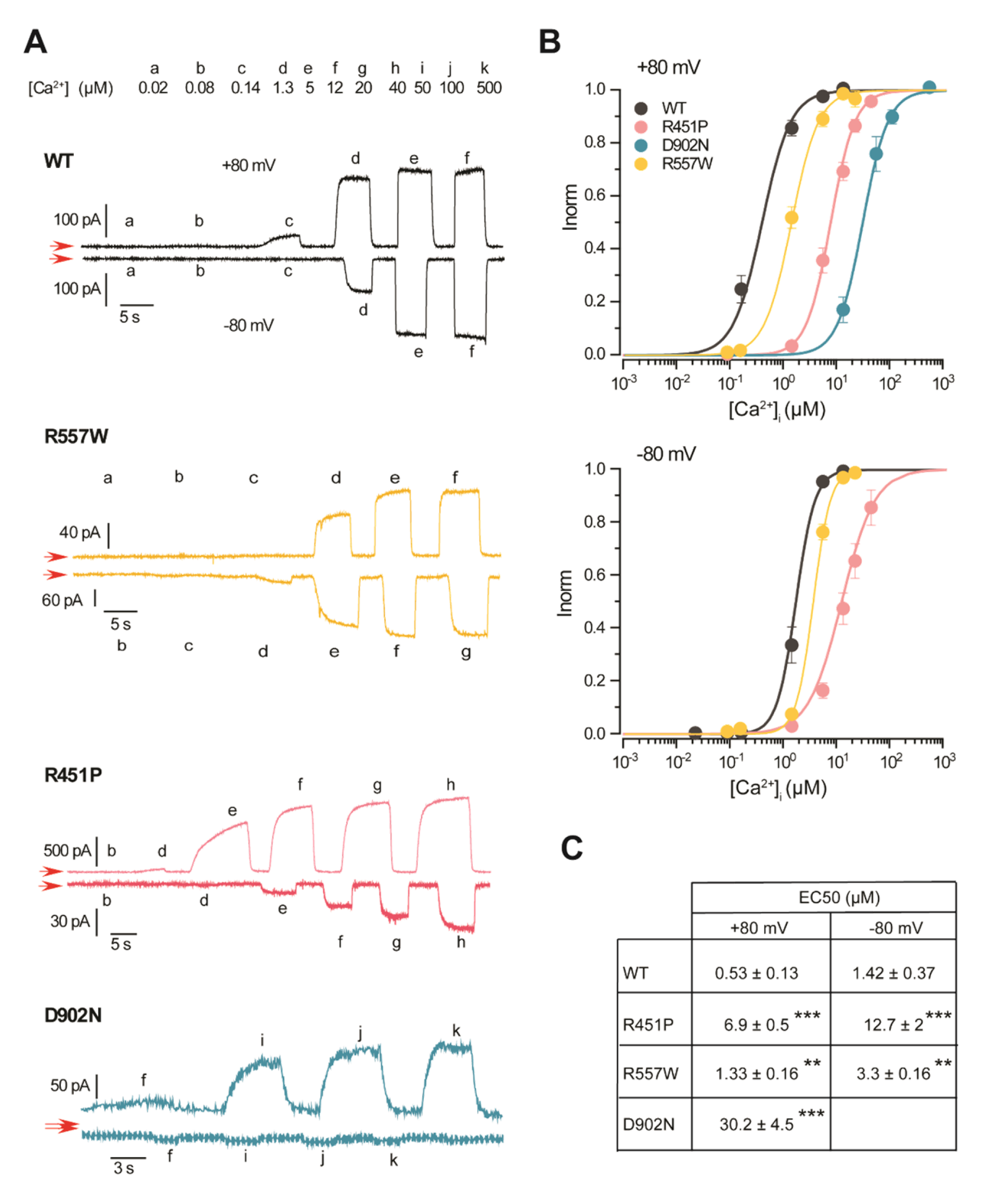
The TMEM16A16A mutants R451P, R557W and D902N have reduced apparent Ca^2+^ sensitivity. **(A)** Representative inside-out patch recordings were obtained from two different patches held at +80 and −80 mV. Patches were exposed to the indicated [Ca^2+^]_i_ (lowercase letters). The red arrows show a 0-nA baseline **(B)** Concentration-response curves to [Ca^2+^]_i_ obtained at +80 mV (upper panel) and −80 mV (lower panel) for TMEM16A WT (black circles, n=6), R451P (pink circles, n=8), R557W (orange circles, n=8) and D902N (cyan circles, n=4). Continuous lines are fits using equation (1) to obtain the EC_50_. At each [Ca^2+^]_i_, the amplitude of I_Cl_ was normalized to the maximum I_Cl_, and the EC_50_ was later estimated by extrapolating the fit to an infinite [Ca^2+^]_i_. **(C)** List of EC_50_ values calculated from the concentration-response analysis. Differences between mutants compared to the WT EC_50_ values were evaluated using the t-student test ***p < 0.001, **p < 0.01.

Altogether, our data demonstrate that R451P, D902N, and K913E point substitutions in the TMEM16A sequence are still functional channels in the plasma membrane. However, these substitutions induced significant changes in their apparent Ca^2+^ sensitivity (R451P and D902N) or produced changes in the opposite direction, mainly eliminating Ca^2+^ dependence (K913E).

### Differential regulation of cell proliferation by low and high Ca^2+^ sensitive TMEM16A mutants

One critical factor in tumorigenesis is the dysregulation of the cell proliferation process. Given that the R451P and D902N mutants displayed low Ca^2+^ sensitivity, whereas K923E have the opposite phenotype, we wondered if these mutants altered cellular proliferation and tumor growth differently. To answer this question, we produced stable transfected OSC-19 cells with the abovementioned mutants, including the D914H, which do not elicit drastic changes in the recorded TMEM16A I_Cl_ at 0.2 µM Ca^2+^. The OSC-19 cell line is derived from human oral squamous cell carcinoma and is a well-established model for subcutaneous cancer to induce tumors in mice (62, 63). OSC-19 cells do not express endogenously high levels of TMEM16A at the protein level. The overexpression of WT, R451P, D902N, K913E and D914H channels was confirmed by Western blotting of plasma membrane enriched samples (Figure 3A).

To determine the function of these mutants in a physiological setting, we measure the ionomycin-induced Cl^-^ fluxes using MQAE, a halide-sensitive dye whose fluorescence is quenched in the presence of Cl^-^. We monitored the changes in MQAE fluorescence intensity upon exposure to an I^-^-containing solution at the same time that we activated TMEM16A by exposing the cells to the Ca^2+^ ionophore ionomycin to increase [Ca^2+^]_i_. Because I^-^ is about 4.5-fold more permeable than Cl^-^ in TMEM16A, its influx at the resting membrane potential quench MQAE, thus giving direct information about TMEM16A activity in intact cells. To compare different experiments using cells expressing other channels, we calculated the relative TMEM16A activity as the changes in MQAE fluorescence divided by the relative plasma membrane expression level.

Under the ionomycin-stimulated condition, we observed an increased anion flux in cells overexpressing TMEM16A relative to parental cells (156.7 ± 23.3 % above its value in parental cells). The observed enhanced anion permeability was sensitive to the TMEM16A channel inhibitor, which reduced the activity to 10.8 ± 0.24% relative to WT TMEM16A expressing cells (Figure 4B, black circles). In contrast, neither ionomycin nor A01 inhibitor affected the mock-transfected control cells (Figure 4B, grey circles).

**FIGURE 4.**
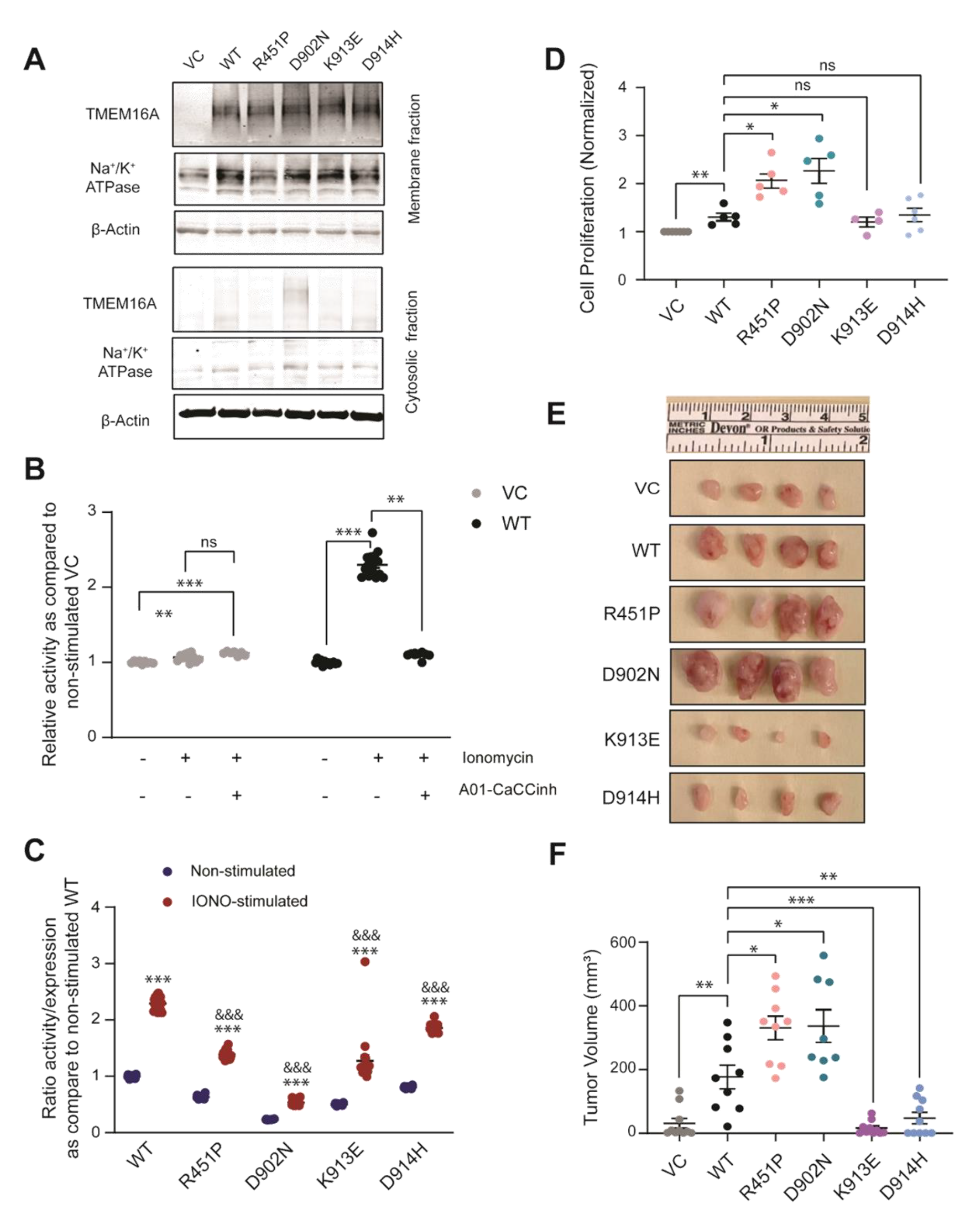
TMEM16A mutations R451P and D902N impact the cell proliferation rate and induce *in vivo* pro-tumorigenic effect. **(A)** TMEM16A overexpression in Oral Squamous Cell Carcinoma cells (OSC-19). Representative Western blot of TMEM16A expression using enriched plasma membrane (upper panel) and cytosolic (lower panel) protein fractions in mock-transfected cells with the empty vector (VC) and in cells expressing WT or mutant channels (R451P, D902N), K913E, and D914H). β-Actin and Na^+^K^+^ ATPase were used as loading control, respectively. **(B)** The activity of TMEM16A was measured using the MQEA quench assay in mock cells, and TMEM16A WT overexpressed stable cell line. TMEM16A activity was stimulated by applying 1 µM ionomycin and inhibited by 20µM A01-CaCCinh 2 h incubation, both added from the extracellular side. Data were normalized to the activity of non-stimulated empty vector cells (VC) (mean ± SEM, n≥3, *** p<0.01 differences with respect to non-stimulated WT cells, ** p<0.01 differences with respect to ionomycin-stimulated WT). **(C)** Activity/expression ratio of WT and mutants TMEM16A in non-stimulated (blue dots) or stimulated with ionomycin (red dots) of ionomycin. The ratio was calculated by dividing the relative MQEA fluorescence (Figure 4B) by the relative membrane expression pattern (Figure 4A). Data were normalized to the value of TMEM16A-WT in non-stimulated conditions. Data were expressed as mean ± SEM, n= 24 from four independent experiments. ***p < 0.001 differences with the respective to non-stimulated condition, ^&&&^p<0.001 differences respect to ionomycin-stimulated WT. **(D)** Cell proliferation rate measurement by WST-1 assay for indicated stable cell lines. Cells were seeded at 4×10^3^ density and analyzed at 72 hr. Data are shown as Mean ± SEM of 4-6 independent experiments *P < 0.05, ** P < 0.01. **(E)** TMEM16A-overexpressing xenografts exhibit differential growth. Bilateral flanks of NOD/SCID mice were injected subcutaneously with OSC-19 TMEM16 WT or the corresponding TMEM16A mutant. Images of harvested OSC-19 tumors are shown for each group. **(F)** Tumor Volume was collected on day 23 post-implantation. Tumor volumes were calculated from data shown in panel E. Differences between groups at ***p < 0.001, **p < 0.01, *p < 0.05. Data represent mean ± SEM of n= 8-10 tumors.

Cells expressing the low Ca^2+^ sensitive mutants R451P and D902N, and those expressing the high Ca^2+^ sensitive K913E channels, showed a significant decrease in the ionomycin-induced activity. Compared to the WT channel, the activity of these mutants decreased by 39.5 ± 1.7, 76.6 ±1.1 and 44.4 ± 9.6 %, respectively (Figure 4C, red circles). Interestingly, the D914H mutant also exhibits a reduction in the relative activity of the channel (19 ± 1.6%), albeit to a lesser extent than the other mutations.

To evaluate if these mutations alter channel activity under low extracellular Ca^2+^ conditions, we measured the MQAE fluorescence in the absence of ionomycin (non-stimulated). We compared the results to those obtained with the WT channel. The relative fluorescence change elicited by the non-stimulated R451P, D902N, K913E and D914H mutants decreased by 36.7 ±1.2, 76.7 ± 0.24, 49.4 ±0.7, and 19.2 ± 0.84 % respectively, relative to the activity generated by the WT TMEM16A. (Figure 4C blue circles). These findings suggest that neither of these mutants exhibits constitutive activity at the resting membrane potential. This idea is supported by the observations that R451P needs an [Ca^2+^]_i_ ≥ 5, µM and D902N is unresponsive to 50 µM Ca^2+^, both at −80 mV and by the strong outward rectification property of all four mutants.

Next, we investigated the effect of TMEM16A R451P, D902N, K913E, and D914H mutations on the proliferation of OSC-19 cancer cells and compared it to the proliferation of cells overexpressing WT TMEM16A. As we expected from previous reports in different cell lines (35, 37, 41, 45, 64-72), WT TMEM16A overexpression induced a 30.4 ± 8% increase in OSC-19 cell proliferation (Figure 4D). In contrast, the expression of the low Ca^2+^-sensitive R451P and D902N mutants led to a much more significant increase in cell proliferation: 107% ± 16 and 127.6 ± 25%, respectively. Surprisingly, the over-expression of the Ca^2+^-independent K913E or the WT-like D914H did not affect OSC-19 cell proliferation (Figure 4D).

To determine the *in vivo* effects of TMEM16A mutations, we used subcutaneous tumor xenografts using OSC-19 implanted in athymic nude mice and measured the tumor volume after 23 days. We terminated the experiment when noticeable differences in tumor size became evident prior to skin ulcerations. Representative pictures are shown in Figure 4E: OSC-19 cells harboring WT TMEM16A produced ∼ five-fold more extensive tumors than did parental controls (Figure 4E). Remarkably, R451P and D902N mutants induced tumors that reached about a ten-fold increase in volume compared to control xenografts (Figures 4D, E). The opposite effect was observed with the K913E mutant channel, dramatically reducing the tumor size (∼ten-fold decrease) relative to tumors induced by OSC-19 TMEM16A WT cells (Figure 4D, E). The volume of K913E tumors was not statistically different from those caused by vector control cells. Similarly, the volume of tumors derived from D914H TMEM16A mutant cells decreased ∼3.5-fold compared with the WT channel. In summary, our results reveal a Cl^-^ flux-independent positive role of low Ca^2+^ sensitive TMEM16A mutants on cell proliferation and tumorigenesis.

### TMEM16A mutants drive cell proliferation and tumor growth through EGFR and AKT signaling pathways

The *in vivo* tumor growth advantage conferred by R451P and D902N mutations and the dominant negative effect of K913E could be driven by changes in signaling pathways associated with cell proliferation, structural changes, or allosteric alterations. Previous data from our laboratory and others highlights the regulatory role of TMEM16A as a key component in Tyrosine Kinase Receptors (EGFR and Her2), Mitogen-activated protein kinase (MAPK) and PI3K/AKT signaling pathways, particularly in cells derived from the head and neck cancer (Bill et al., 2015; Britschgi et al., 2013; Crottes et al., 2019; Duvvuri et al., 2012). Given that over-activation of these cascades is associated with cell survival, inducing cell cycle deregulation, altering the processes of proliferation, differentiation, migration, and angiogenesis (Wee and Wang, 2017), we tested the effect of WT and R451P, D902N, K913E mutants on the phosphorylation state of EGFR, AKT and ERK as an indication of the cell signaling pathway activation.

In line with this idea, we found that overexpression of WT TMEM16A increased EGFR (Y1060), AKT (S473), and ERK1/2 (T202/Y204) phosphorylation by 27.7 ± 11.7 %, 47.6 ± 15.1% and 21.6 ± 1%, respectively. Cells overexpressing the R451P mutant increased the AKT S473 phosphorylation to 126% ± 23.4 %.

Nevertheless, the activation of EGFR (21.1 ± 21.8%) and ERK1/2 (15.1 ± 4.9) remains unaltered. Contrary to its pro-tumorigenic effect, neither AKT nor ERK1/2 exhibited over-phosphorylation in cells expressing the D902N mutant. No statistical differences were observed between cells expressing D902N and empty vector (Figure 5A, B). Remarkably, the overexpression of D902N TMEM16A mutant induced a two-fold increase in EGFR phosphorylation (increase to 170.7 ± 24.3 %) (Figure 5A, B), suggesting that this version of TMEM16A may promote cell growth by switching signaling control from AKT to EGFR. Interestingly, the K913E mutant that reduced cell proliferation and tumor growth exhibited a 55.7 ± 15.8% increase in the activation of EGFR. However, this mutant also showed decreased phosphorylation levels of AKT and ERK1/2, ultimately reaching similar levels to the endogenous EGFR activity in OSC-19 cells.

**FIGURE 5.**
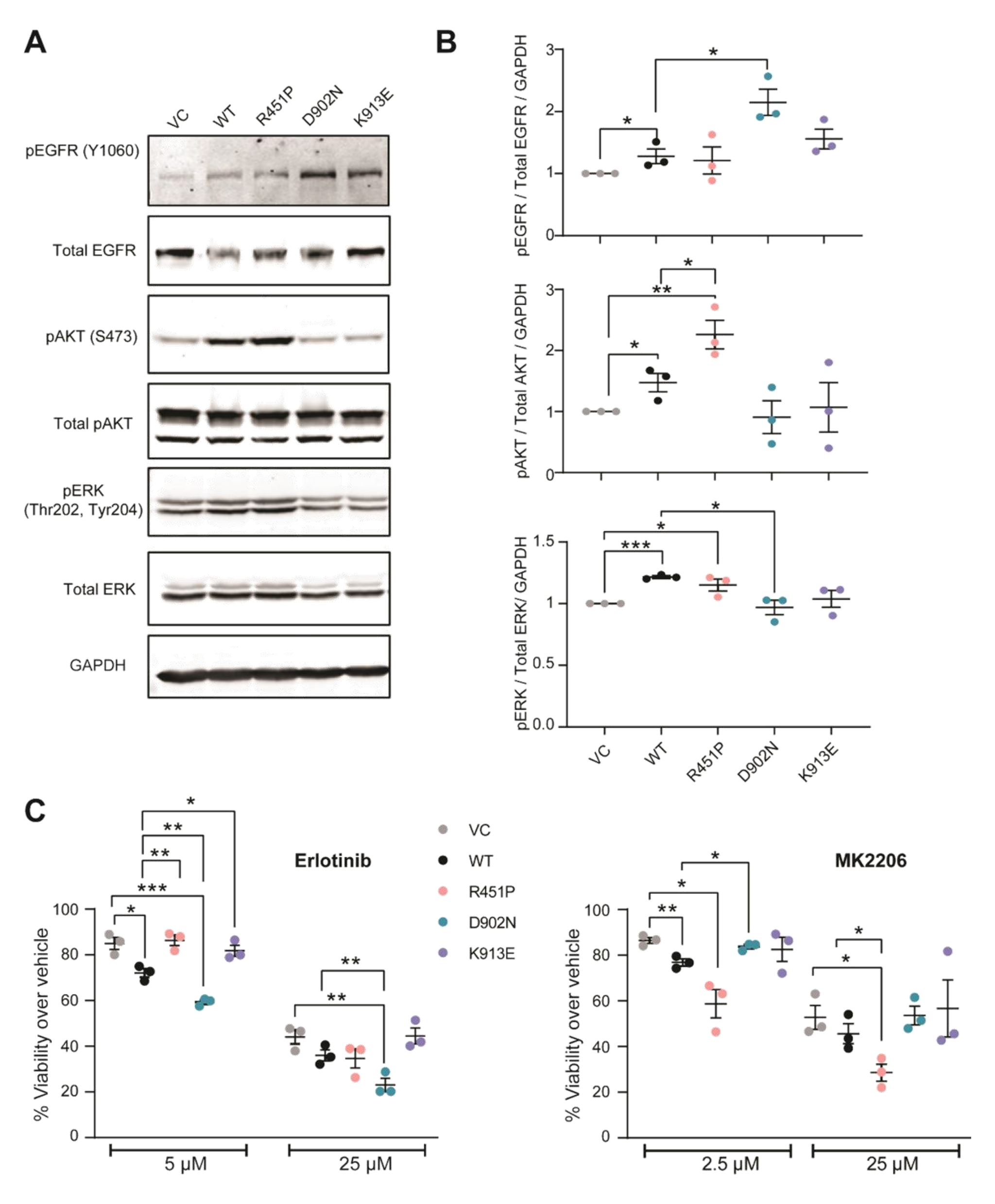
The TMEM16A R451P and D902N mutants have a pro-oncogenic phenotype that correlates with the functional activation of AKT and EGF receptor. **(A)** Immunoblotting analysis of OSC-19 cells over-expressing WT, R451P, D902N, and K913E channels. Proteins extracted from three independent sets of each cell type were subjected to Western blot analysis using the antibodies phospho-EGFR (Y1060), phospho-AKT (S473), Total AKT, phospho-ERK1/2 (T202/Y204), and Total ERK1/2. GAPDH was used as a housekeeping protein. **(B)** Optical density ratio phospho-protein/Total protein/GAPDH in empty vector (VC), WT, and mutant channels. Values expressed as mean ± SEM, ***p < 0.001, *p < 0.05, t-test, n=3 independent experiments. (**C**) Cell proliferation of cells treated with the indicated concentration of EGFR (Erlotinib, left panel) and AKT (MK2206, right panel) inhibitors 72h. Values were normalized to the respective vehicle-treated cells and are presented as the mean ± SEM of three-four independent experiments. (*p < 0.05; **p < 0.01; p < 0.001***).

To further test the role of EGFR or AKT activation in OSC-19 cell proliferation, we measured cell viability in the presence of low and high doses of EGFR and AKT inhibitors Erlotinib and MK2206. We find that Erlotinib interferes with TMEM16A-induced proliferation in OSC-19 cells. Figure 5C shows a 15 ± 2.6 % of survival decrease of control cells treated with 5 µM Erlotinib compared to the vehicle (DMSO) treated cells. WT and D902N cells are significantly more sensitive to 5 µM Erlotinib, which caused a 27.9 ± 1.9, and 40.7 ± 0.9 % of cell survival decrease, respectively. Interestingly, despite the increased phosphorylation levels of EGFR in the K913E mutant, no significant changes in inhibitor sensitivity were observed (18.2 ± 2.4%). Similar observations in the cell survival sensitivity to EGFR inhibitor were made at high concentrations of Erlotinib, as shown in Figure 5C. These findings strongly suggest that the pro-oncogenic capacity of the D902N mutant is related to the over-activation of EGFR signaling. On the other hand, the K913E mutant may engage distinct signaling cascades that bypass the effects of EGFR activation, thereby maintaining a slow proliferation capacity.

To probe if the AKT signaling pathway is involved in the pro-oncogenic effect of the R451P mutant, we used MK2206, an inhibitor of AKT. Treatment with low doses of the inhibitor resulted in a notable reduction in cell proliferation of R451P cells (23.2 ± 1.5% decrease), significantly different from the reduction in cell survival observed in empty vector cells (13.5 ± 0.26%). No statistical differences were observed in the viability of cells expressing the mutant D902N (16.2 ± 0.9% decrease) or K913E (17.4 ± 5.3% decrease) compared to control cells. However, mutant R451P exhibited a pronounced increase in MK2206 sensitivity, demonstrating a substantial reduction of 41.3 ± 6.2% in cell survival upon treatment. Increasing the MK2206 concentration to 25 µM enhanced inhibition, as evidenced by a 71.4 ± 3.7% decrease in cell survival in R451P mutant cells compared to WT TMEM16A (54.4 ± 4.4%) (Figure 5D). This finding provides evidence that the overactivation of the AKT pathway by the mutant R451P cells is likely responsible for the increased cell proliferation and tumorigenesis. The increased sensitivity of the mutant cells to MK2206 supports the notion that the aberrant activation of the AKT pathway plays a crucial role in promoting enhanced growth and tumor formation by cells expressing these mutant channels.

### Structural changes may underline the cell proliferative effect of R451P and D902N mutant channels

The binding of Ca^2+^ is obligatory for channel activation and subsequent ion conduction. A salient characteristic of R451P and D902N is their low Ca^2+^ sensitivity, which most likely precludes their ion conduction function under physiological conditions. However, their low Ca^2+^ sensitivity did not hamper their cell proliferative and tumorigenic phenotype. This result suggests that the Ca^2+^-unbound state of the channel is most likely participating in proliferation, tumor growth and cell signaling.

To test this hypothesis, we replaced the side chains of R451 and D902 with charge-neutralizing (R451Q) and charge-conserving (D902E) amino acids and tested their Ca^2+^ dependence. We find that the R451Q mutant rescued the Cl^-^ current activated by 0.2 µM Ca^2+^ and completely restored the Ca^2+^ sensitivity to a value (EC_50_= 0.47 ± 0.06 µM and 1.67 ± 0.07 µM at +80 and −80 mV respectively) that is not statistically different from that determined for WT TMEM16A (Figure 6C, D). Similarly, the D902E mutation rescues both the I_Cl_ current (Figure 6A, B) and its apparent Ca^2+^ sensitivity (EC_50_ = 0.58 ± 0.03 µM at +80 mV and 1.78 ± 0.20 µM at −80 mV; Figure 6C, D).

**FIGURE 6.**
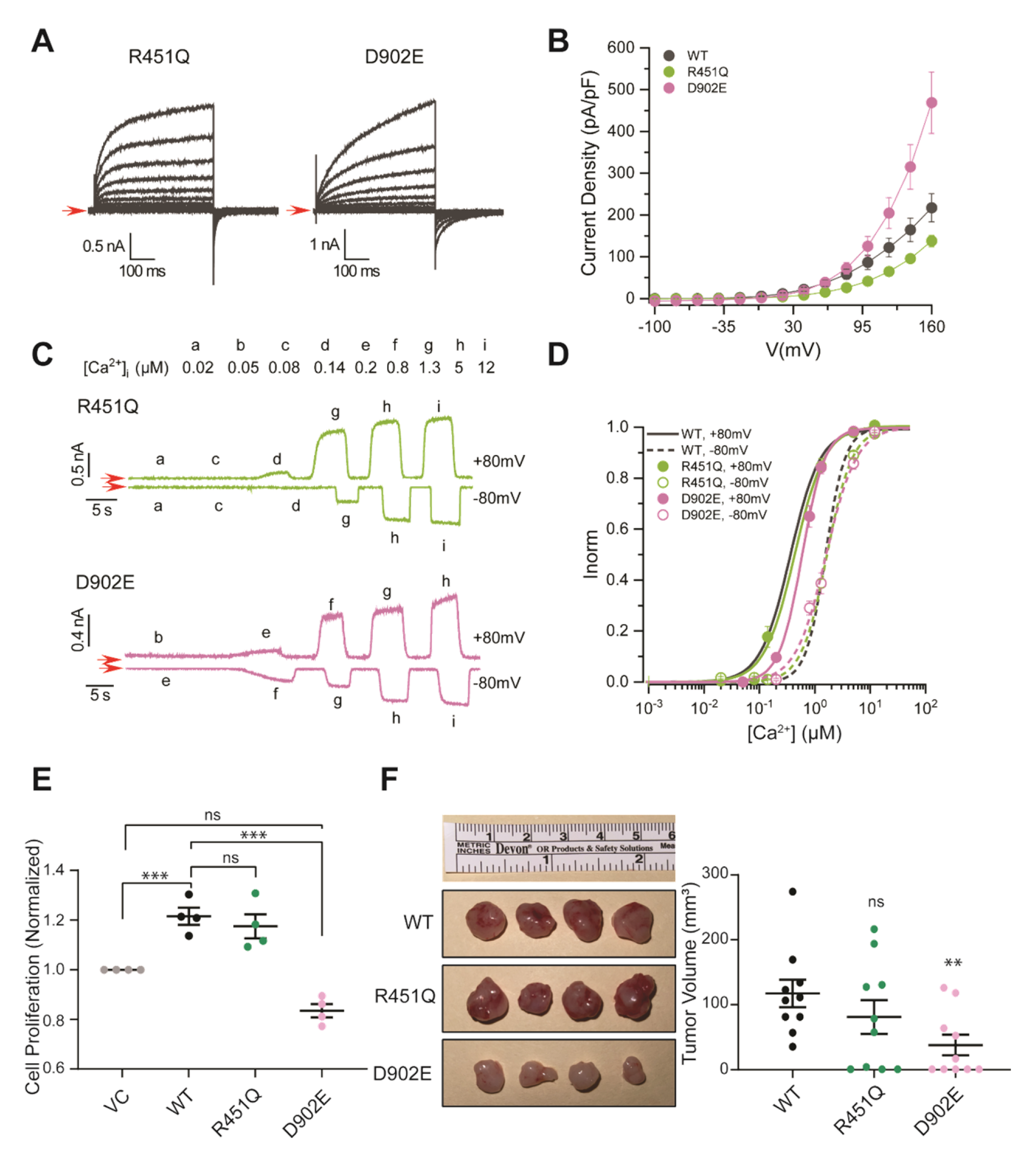
Recovery of channel function, cell proliferation and tumor growth by mutants with conservative Amino Acid Side Chains at position 451 and 902 of TMEM16A. **(A)** Representative whole-cell I_Cl_ recordings from HEK-293T cells expressing TMEM16A R451Q and D902E. Recordings were made using a pipette solution containing [Ca^2+^]_i_= 0.2 µM. The stimulation protocol is shown in Figure 2B. The Red arrows indicate a 0-nA baseline **(B)** Mean steady-state whole-cell current density versus V_m_ relationships constructed using the current measured at the end of the varying step in the activation protocol. TMEM16A WT (black circles n=8), R451Q (green circles n=6), and D902E (pink circles n=10). **(C)** Representative Inside-out patch recordings were obtained at +80 and −80 mV from two different patches excised from cells expressing R451Q (green) or D902E (pink). Patches were exposed to the indicated [Ca^2+^]_i_ (lowercase letters). **(D)** Concentration-response curves to [Ca^2+^]_i_ were obtained at +80 mV with TMEM16A R451Q (green-filled circles, n=8) and D902E (pink-filled circles, n= 9). Curves obtained at −80 mV are in empty circles, R451Q n=8, D902E n=8. Continuous and dashed lines are fits using equation (1). At each Ca^2+^ concentration, the amplitude of I_Cl_ was normalized to the maximum level; the EC_50_ was later estimated by extrapolating the fit to an infinite [Ca^2+^]_i_. **(E)** Cell proliferation in control, R451Q and D902E cells after 72 h. Data presented as mean ± SEM (n=4, *p < 0.05). **(F)** R451Q and D902E TMEM16A-overexpressing xenografts do not exert a pro-tumorigenic effect. Differences between R451Q and D902E cells’ tumor volume compared to WT TMEM16A cells were evaluated using the t-student test **p < 0.01. Data represent mean ± SEM of n=10 tumors.

Because R451Q and D902E are WT-like channels, we wanted to know whether cell proliferation and tumor growth also reverted to values like those produced by the WT channel shown in Figure 4D. Amazingly, TMEM16A R451Q overexpression increased *in vitro* OSC-19 proliferation by 17.5 ± 4.8%. This proliferation increase was significantly smaller than the proliferation induced by the R451P mutant but not statistically different from that produced by the WT channel (21.5 ± 3.4%). Surprisingly, TMEM16A D902E mutant showed a dominant-negative effect on cell proliferation. This mutant abrogated the increase in cell viability produced by overexpressing the TMEM16A WT and D902N mutant (Fig. 6E).

Finally, we determine the effects of these WT-like mutants on tumor growth. In agreement with the *in vitro* assays, the results revealed that tumor volumes induced by the R451Q mutant were not statistically different to those produced by the WT (81 ± 24.8 mm^3^ *vs* 117.4 ± 20.2 mm^3^, respectively). Similarly, the tumor volume (38 ± 15 mm^3^) induced by the overexpression of the D902E mutant decreased by nearly three-fold compared to tumors induced by WT TMEM16A cells (Figure 6F).

These data suggest that alterations in the channel structure associated with the proline insertion at position 451 and charge neutralization in the D902 residue are responsible for the tumorigenic effect.

## DISCUSSION

A growing amount of evidence supports the involvement of WT TMEM16A in cancer development (29-31); however, the role of mutations in the oncogenic process remains unknown. More than four hundred mutations in TMEM16A have been reported on the cBioPortal for Cancer Genetics, a dedicated platform to determine protein mutations in different cancer samples. The analysis of single, double, or triple mutations in specific cancer represents a paramount work beyond the scope of this work. This is why in this work, for the first time, we investigated the functional and oncogenic implications of nine-point mutations identified in the channel, which were previously reported in human cancerous tissues, including Head and Neck Squamous Cell Carcinoma, Cutaneous Melanoma, Invasive Breast Carcinoma, Stomach Adenocarcinoma, Thymoma, Uterine Endometrioid Carcinoma, Prostate Adenocarcinoma, Glioblastoma Multiforme, and Lung Adenocarcinoma. The strategic location of the mutants we studied (R451P, R455Q, M546I, R557W, F604L, D902N, K913E, D914H, and Q917K), at the extracellular domain and at the TM2-TM10 joint surrounding the third Ca^2+^ binding site that allosterically regulates the canonical site Ca^2+^ binding site 1 (73), led us to hypothesize that alterations in the Ca^2+^ sensitivity maybe underlie cell proliferation and cancer. In agreement, we found that the R451P and D902N mutations have decreased Ca^2+^ sensitivity, activate the AKT and EGFR signaling pathways, and are oncogenic.

The oncogenic phenotype of R451P and D902N is notable. R451P activates the AKT pathway and increases OSC-19 cell proliferation. These proliferative effects of R451P were reversed by the AKT inhibitor MK2206 and by mutating R451 to Q instead of P, thus clearly demonstrating an activating role of this mutation on the AKT signaling pathway. Furthermore, xenografts carrying the R451P mutant channels grow significantly more than those produced by the WT channel. On the other hand, the signaling pathway activated by the D902N TMEM16A mutant was different from that induced by WT and R451P channels. This mutant promotes cell proliferation and increases the EGFR but not AKT phosphorylation. Nevertheless, the charge conservation mutant D902E reduced cell proliferation to lower levels than those observed in cells expressing the WT channels. The increased AKT/EGFR activation observed in these mutants provides a potential mechanistic explanation for its oncogenic properties, highlighting the crucial role of these signaling pathways’ dysregulation in driving tumor progression. Our data suggest that TMEM16A likely leverages several ways to promote tumor growth and that each path requires a specific aspect of TMEM16A function. Therefore, treating TMEM16A overexpressing tumors with channel blockers might fail not just because they are not specific enough but because they do not distinguish between TMEM16A mutants and, therefore, might not inhibit every aspect of TME16A function that drives cell growth and proliferation (74). Interestingly, the alignment of TMD6 and TMD10 of TMEM16A shows that R451 and D902 residues are conserved between species. Residue R451 is also conserved in other TMEM16 family members, including the Ca^2+^-activated Cl^-^ channel TMEM16B and the scramblase TMEM16F (Supplemental Figure 2). Whether this indicates a conserved phenotype is unknown.

The overexpression of TMEM16A in cancer cells could enhance the Cl^-^ efflux, potentially altering cellular signaling and increasing proliferation. However, our data do not support this hypothesis. Firstly, the proliferation and tumor growth experiments were done without stimuli that would increase the intracellular Ca^2+^. Under this condition, the contribution of the Cl^-^ efflux would be minimal. Second, the patch clamp data, which measure the activity of channels in the plasma membrane, showed that R451P and D902N mutants produced about 4.5-fold less current than WT, even though they were recorded at significantly higher Ca^2+^ concentrations. Note that the 1.5 µM Ca^2+^ used in the R451P recordings is slightly below its EC_50_ (6.9 µM), whereas D902N was activated in the presence of 50 µM Ca^2+^, well above its EC_50_ (30.2 µM). The strong rectification of the currents is translated into a tiny current in the negative range of Vm near those values encountered under physiological conditions. In addition, these oncogenic mutants displayed a relatively low Ca^2+^ sensitivity, implying that their activation in cancer cells at the negative resting membrane potential is poor. Thus, it is expected that such a tiny current would not deplete the intracellular Cl^-^ content by any significant amount. The functional analysis of TMEM16A by the halide-sensitive sensor shows a decrease in the Cl^-^ flux induced by ionomycin in intact cells expressing R451P and D902N thus supporting this idea. Despite these properties, both R451P and D902N mutants enhanced cell proliferation and tumor growth. Therefore, we propose that the Cl^-^ efflux via TMEM16A channels does not play a role in determining their oncogenic phenotype. Instead, TMEM16A mutants differentially activate the signaling pathways AKT and EGFR to promote cell growth and proliferation; the fact that such activation was differential illustrates the complexity of signaling pathways set on by TMEM16A. This idea is supported by the R451N and D902E mutants that showed WT-like behavior, and the cell proliferation and tumor size were reduced to WT levels or even lower.

However, an opposite result is illustrated by the K913E mutant. In this case, Vm can activate the mutated channel in the absence of intracellular Ca^2+,^ and the current *vs* voltage curve shows strong outward rectification, which generates a very small current at negative potentials (Figure 2B), not enough to disturb the intracellular Cl^-^ content. MQEA assays confirm a minor alteration in Cl^-^ flux induced by the K913E mutant, both in the absence of stimulation and upon elevating the Ca^2+^ using ionomycin. Remarkably, xenografts carrying the K913E channel have dramatically decreased tumorigenic capacity. This effect could be attributed to the reduction in the activation levels of AKT and ERK1/2, which aligns with the endogenous levels observed in OSC-19 cells.

In summary, we show that the Cl^-^ flux via TMEM16A channels is dispensable in tumorigenesis. In addition, our work shows that specific cancer-associated TMEM16A mutations drive the oncogenic process. This idea is nicely illustrated by the residues R451 and D906 that enhance the cel’s oncogenic capability when they are mutated to P and N, but not when they are mutated to Q and E, respectively. Also, our work revealed that different mutants activate different proliferative signaling pathways, which emphasizes the channe’s multifaceted role in cell proliferation and tumor development.

## Supporting information

curated list

## Author Contributions

Concept Design: UD, SC-R

Acquisition of Data: SC-R AG, RG, JJJP, JP, MRB, AYK

Analysis and interpretation of data: SC-R, JJJP, JA, KK, UD

Writing and review of the manuscript: SC-R, KK, UD, JA

Provided microscope facilities: G.R.V.H.

## Competing Interest

The authors declare no conflict of interest.

Data Availability Statement

The data supporting this study’s findings are available on request from the authors.

## Abbreviations

ANO1: Anoctamin-1
CaCC: Calcium-activated chloride channel
[Ca^2+^]_i_: Intracellular calcium concentration
CAMKII: Ca^2+^/Calmodulin-dependent protein kinase II
EGFR: Epidermal Growth Factor Receptor
ERK1/2: p44/42 Mitogen-activated protein kinase
I_Cl_: Chloride current
IONO: Ionomycin
MAPK: Mitogen-activated protein kinase
NFκB: Nuclear factor κB
OSC-19: Oral Squamous Cell Carcinoma cells
PtdIns(4,5)P2: Phosphatidylinositol 4,5-bisphosphate
TMD: Trans-membrane Domain
TMEM16A: Transmembrane protein 16A
T_on_: Activation time constant
T_off_: Deactivation time constant
Vm: Transmembrane Voltage

## ACKNOWLEDGEMENTS AND FUNDING

This work was supported in part by grants from the Department of Veterans Affairs (IO1-002345) and the NIH (RO1-DE028343 (to U.D.). FORDECYT-PRONACES 1308052 2020 from CONACyT, Mexico (to JA).

The contents do not represent the views of the Department of Veterans Affairs, or the United States Government.

We wish to acknowledge the technical assistance of Carmen Y. Hernández-Carballo.

## METHODS

### Cell lines

Site-directed mutagenesis was performed on hTMEM16A (GenBank NM_001378095.2) by using Quick-change mutagenesis kit (Aligent, Santa Clara, CA, USA) according to the manufacturer’s protocol and cloned in pBabe-puromycin retroviral vector as a template. All mutations were confirmed by Sanger sequencing (GeneWiz). TMEM16A WT and mutants R451P, R455Q, M546I, R557W, F604L, D902N, K913E, D914H, and Q917K were subcloned into PCMV-AC GFP vector for electrophysiological recordings in HEK293 cells. HEK293 cells were maintained in Dulbecco’s modified Eagle’s medium (DMEM) supplemented with 10% fetal bovine serum and 1% Penicillin/Streptomycin at 37°C in a 95% O_2_–5% CO_2_ atmosphere.

OSC-19 cells were maintained in high glucose Dulbecco’s modified Eagle’s medium supplemented with 10% FBS and 1% Penicillin/Streptomycin mixture. Platinum Retroviral Packaging Cell Line (Plat-A) was cultured in Dulbecco’s modified Eagle’s medium containing 10% FBS, Puromycin 1 μg/ml, and Blasticidin 10 μg/ml. Retrovirus particles were collected from Plat-A cells transiently transfected with 8 μg of the respective TMEM16A pBabe constructs and were used to transduce OSC-19 cells. Retroviral transduction was made by using Polybrene 10 μg/ml and 1 ml virus stock. For antibiotic selection, 1 μg/ml puromycin was added to the medium. Cells were incubated in the presence of puromycin for 10 days to obtain a pooled population of cells.

All cell lines were mycoplasma-free and authenticated using human cancer cell line STR profiles. Stable cell lines were used for eight passages and then discarded.

### Electrophysiological recordings

According to the manufacturer’s instructions, HEK293 cells were transiently transfected with 1 µg of each cDNA using Lipofectamine 2000 (Invitrogen, Carlsbad CA, USA). After 12 h of transfection, cells were plated onto 5 mm glass coverslips pre-coated with 0.1 mg ml−1 poly-L-lysine and allowed to attach for at least 4 h before recordings of the Cl^-^ currents (I_Cl_). The experiments were conducted at room temperature (21°C) using the whole-cell patch-clamp technique with an Axopatch 200B amplifier (Molecular Devices, Sunnyvale, CA, USA). The voltage protocol consisted of a holding potential of −60 mV followed by voltage steps from −100 to +160 mV in 20 mV increments lasting 0.5 s, then a repolarizing step to −100 mV. The time interval between pulses was 7 s. Data acquisition and generation of voltage protocols were carried out using a Digidata 1550 interface (Molecular Devices). Whole-cell currents were filtered at 5 kHz and digitized at 10 kHz using the software pCLAMP 10 (Molecular Devices). Micropipettes were pulled from Corning 8161 capillary glass tubes (Warner Instruments, LLC, Hamden, CT, USA) using a PP830 electrode puller (Narishige, Tokyo, Japan). Pipette resistances ranged from 3 to 6 MΩ.

The 0.2 µM [Ca^2+^]_i_ intracellular solution contained (mM) 30 NMDG-Cl, 20 HEPES, 5.2 CaCl_2_, 25.2 EGTA, 85 D-mannitol pH was adjusted to 7.3 with Tris, osmolality 290–300 mosmol kg^−1^. The final free Ca^2+^ concentration was estimated by the MAXCHELATOR program (maxchelator.stanford.edu). Cells were bathed in a standard external solution that contained (in mM): 140 NMDG-Cl, 1.5 CaCl_2_, 10 Hepes, and 100 D-mannitol, pH adjusted to 7.3 with Tris, 380-400 mosmol kg^−^1. The osmolality of all solutions was measured using the vapor pressure point method (VAPRO, Wescor Inc., South Logan, UT, USA).

Data were analyzed and plotted using pCLAMP 10 (Molecular Devices) and Origin 10 (Origin Lab, Northampton, MA, USA). The experimental data are presented as means ± SEM. I_Cl_ *vs* V_m_ curves were constructed using the I_Cl_ values at the end of each depolarizing pulse normalized to the I_Cl_ magnitude recorded at +160 mV in the same cell.

To test the channel activation by intracellular Ca^2+^, inside-out patches excised from cells expressing the WT or the indicated mutants were held at +80 or −80 mV and exposed to [Ca^2+^]_i_ ranging from 0.01 to 500 μM (Supplementary Table 1). To check for changes in seal resistance, each test [Ca^2+^]_i_ response was preceded by exposure to a control Ca^2+^-free solution containing 25.2 mM EGTA. The current from the inside-out patches were filtered at 1 kHz and digitized at 10 kHz. Concentration dose–response curves were constructed after normalizing the individual responses to the maximum current obtained with the highest [Ca^2+^]_i_ in the same patch and then pooling the normalized data. The resulting curves were fit to

Hill’s equation:

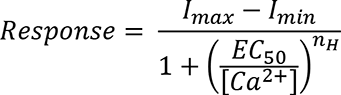

where *I_max_* − *I_min_* are the maximum and minimum current responses, respectively, EC_50_ the [Ca^2+^] to have the 50% of the response and n_H_ is the Hill coefficient.

### Animal studies

Fifteen NOD-SCID gamma female mice (4 weeks old) were implanted subcutaneously with 3×10^6^ OSC-19 cells in each flank. Mice (n=6-7) were randomized in groups: vehicle, TMEM16A WT and mutants R451P, D902N, and K913E, D914H, R451Q and D902E according to the experiment. The Volume of the tumors was measured at the end of 23 days. Termination of the experiment was required to avoid mouse skin ulceration. Animals were handled and euthanized following IACUC guidelines.

### Confocal Microscopy

HEK293 cells were plated on 35 mm tissue culture dishes with 20 mm Number 1.5 cover glass apertures (CellVis) previously coated with 5 μg fibronectin (Life Technologies 428 33016-015). Cells were transiently transfected with 100 ng of empty vector, TMEM16A WT or the corresponding TMEM16A mutant channel. Twenty-four hours after transfection, cells were imaged. The standard growth medium was replaced with Fluorobrite DMEM (Thermo Fisher; A1896702) supplemented with 10% heat-inactivated FBS, 25 mM Hepes (pH 7.4), 0.1% chemically defined lipid supplement, and 2 mM GlutaMAX (Thermo Fisher; 35050061).

Cells were imaged on a confocal Nikon A1R equipped with a 100X 1.45 NA plan-apochromatic objective. A resonant mode was used to capture all images. Excitation of fluorophores was accomplished with a 488 nm laser mounted in a dual fiber-coupled LUN-V laser launch. Emission was collected with a green filter with a 500–550 nm bandpass. Confocal pinhole size was defined as 1.2 Airy disc size at 488 nm. Nikon Elements software was used to acquire images.

### Western Blot analysis

Cell samples were homogenized in ice-cold lysis buffer (1 M Tris-HCl pH 7.6, 0.5 M EDTA, 5 M NaCl, 0.1 M Na_4_P_2_O_7_, 0.5 M NaF, 0.05 Na_3_VO_4_, 1% Triton X-100) containing protease and phosphatase inhibitor cocktail (Thermo Fisher Scientifics, Rockford, lL 61105 USA). The protein was separated by SDS/PAGE gel and transferred to nitrocellulose membranes. Membranes were blocked for 1 h at room temperature and incubated with primary antibodies overnight at 4°C. Proteins were detected using Anti-TMEM16A from Thermo Fisher (#MA5-16358, Thermo Fisher Scientific, Waltham, MA); pEGFR (Y1068, #3777) pAKT (S 473, #9271 and p44/42 MAPK (ERK1/2) (T202/Y204, #4370) were purchased from Cell Signaling Technology (Danvers, MA), β- actin was used as a housekeeping proteinand the antibody was bought from EMD Millipore (#MAB1501 now part of Millipore Sigma, Burlington, MA). After secondary antibody incubation, Imaging in the LiCor Odyssey system or standard camera luminescence. Densitometry analysis was conducted using signal quantification software provided in the LiCor Odyssey system. Cropped images are presented for conciseness, and the brightness or contrast was adjusted uniformly to the entire image.

### Plasma membrane enrichment

Plasma membrane protein enrichment was carried out using the Thermo Mem-Per Plus kit (Thermo Fisher Scientific, Waltham, MA) following the manufacturer’s instructions. In summary, 15×10^6^ cells were harvested and subjected to two washes with Cell Wash Solution before being treated with permeabilization solution containing protease and phosphatase inhibitor cocktail (Thermo Fisher Scientifics, Rockford, lL 61105 USA). Cells were vortex briefly and incubated for 10 minutes at 4°C with constant mixing. After centrifugation of 15 minutes at 16,000 x *g,* we carefully removed the supernatant containing cytosolic proteins. Pellet was resuspended in Solubilization Buffer and incubated at 4°C with constant mixing for 30 minutes. The supernatant containing solubilized membrane-associated proteins was recovered by centrifugation at 16,000 x *g* for 15 minutes. Protein quantification and analysis of both fractions were performed using the Western blot technique. Na^+^/K^+^ ATPase from Cell Signaling Technology (#3010S, Danvers, MA) and β- actin from EMD Millipore (now part of Millipore Sigma, Burlington, MA) were used as housekeeping protein, and β-actin antibodies (Cell Signaling) were used as a control of enrichment of membrane and cytoplasmic fractions respectively.

### Modeling protein structure

The hTMEM16A structure model (Q5XXA5) was downloaded from AlphaFold Protein Structure Database (75, 76). Structure regions with a confidence score (pLDDT) < 50 were discarded. α2 and α10 helixes have a pLDDT > 83. Ca^2+^ ions and dimer were placed and built, aligning the TMEM16A model and the TMEM16A cryo-structure (7ZK3,(77)) using ChimeraX.

### Viability assay

A cell-viability assay was performed according to the manufacturer’s protocol by using 2-(4-iodophenyl)-3- (4-nitrophenyl)-5-(2,4-disulfophenyl)-2H-tetrazolium, monosodium salt (WST-1) (Takara BioInc, Clontech Laboratories, Inc.). 4 × 10^3^ cells/ml was cultured in 5 replicates in 96-well plates. Cells were treated with Erlotinib 5 or 25 μM or MK2206, 2.5 or 25 μM in DMEM FBS-free media 24 h after seeding. The treatment was carried out for 48 hr at 37^0^ C, followed by measurements of viability; absorbance was read at 450 nm in a microplate reader from BioTek Instruments, Inc. Relative proliferation readings from the treated groups were normalized to the readings from the vehicle-treated cells.

### Halide fluxes measurement

For measurements of the ionomycin-induced halide flux from cells overexpressing the TMEM16A channel, 25×10^3^ OSC-19 cells were grown in 96-well plates. Cells were loaded overnight with 5 mM MQAE (Invitrogen, Carlsbad CA, USA) in 100 μl/well of DMEM FBS 10%. Cells were washed 2 times with 80 μl of buffer solution (in mM: 135 NaCl, 2 KCl, 1 CaCl2, 1 MgCl_2_, 10 glucose and 5 HEPES). Measurements were taken at 2-min intervals using an excitation wavelength of 350 nm and an emission wavelength of 460 nm for 6 min to reach a baseline fluorescence in a BioTek Instruments, Inc microplate reader. A total of 20 μl of assay solution (in mM: 350 NaI, 2 KCl, 1 CaCl_2_, 1 MgCl_2_, 10 glucose, 5 HEPES, pH 7.4) was added in the absence or presence of 5 µM ionomycin (final concentration 1 μM). Fluorescence was continuously recorded for 10 min. Halide flux was evaluated in the WT channel and the mutant variants by comparing the fold change in the fluorescence following exposure to the assay solution, which was subsequently normalized against the relative membrane protein level of the channel.

### Statistical analysis

Data analysis and generation of plots were performed with Prism 9.0 (GraphPad) and Origin 10 (OriginLab, Northampton, MA). Results are presented as the mean ± standard error of the mean (SEM). The statistical significance was determined using a Student’s t-test, with the P-value reported exactly or one-way ANOVA with post hoc Bonferroni test for correction of multiple comparisons.

Values were considered significantly different if p<0.05.

