## Supplemental Figure 1 for "Missense mutations in the calcium-activated chloride channel TMEM16A promote tumor growth by activating oncogenic signaling in Human Cancer"

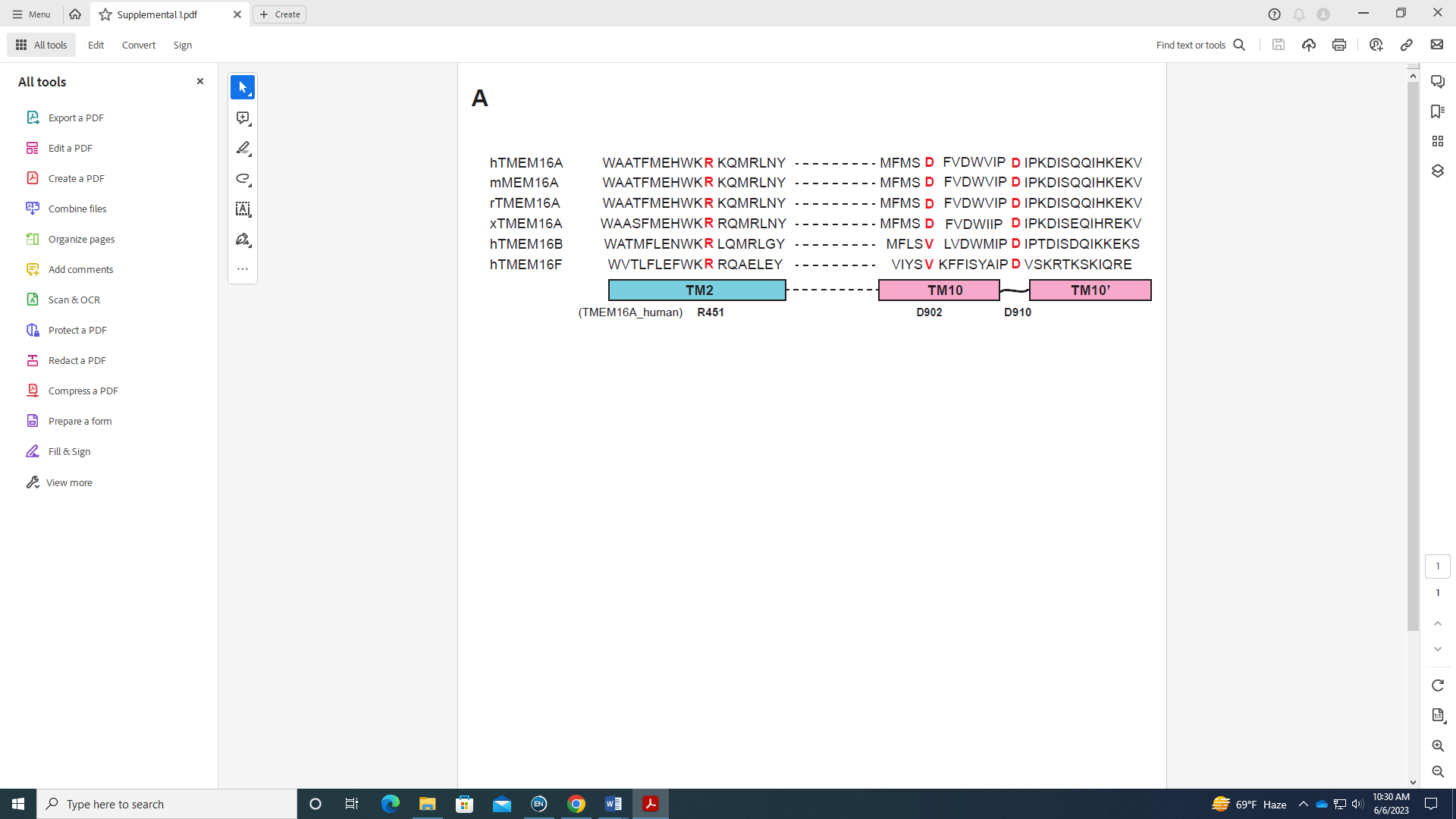

**Supplemental Figure 1**. Sequence alignment of the transmembrane region 2 and 10 (TM2, TM10) of human TMEM16A (hTMEM16A), mouse TMEM16A (mTMEM16A), rat TMEM16A (rTMEM16A), *Xenopus* TMEM16A (xTMEM16A), human TMEM16B (hTMEM16B) and human TMEM16F (hTMEM16F). Residues R451, D902 and D910 are highly conserved among distinct species and TMEM16 family members (shown in red).
