## Supplemental Table 1 for "Missense mutations in the calcium-activated chloride channel TMEM16A promote tumor growth by activating oncogenic signaling in Human Cancer"

**Supplemental Table1. Composition of intracellular solutions used in inside-out patch-clamp experiments.**

|  | **Free [Ca^2+^] (µM)** | | | | | | | | | | | | |
| --- | --- | --- | --- | --- | --- | --- | --- | --- | --- | --- | --- | --- | --- |
| **Compound**  **(mM)** | **0** | **0.01** | **0.02** | **0.05** | **0.08** | **1.3** | **5** | **12** | **20** | **40** | **50** | **100** | **500** |
| **NMDG** | 51.2 | 51.2 | 51.2 | 5.2 | 51.2 | 51.2 | 51.2 | 51.2 | 51.2 | 51.2 | 51.2 | 51.2 | 51.2 |
| **HCl** | 51.2 | 36.5 | 40.72 | 33.68 | 28.2 | 38.6 | 23 | 13.2 | 9.2 | 5.4 | 4.6 | 2.8 | 0.6 |
| **CaCl2** | - | 7.35 | 5.241 | 8.76 | 11.5 | 6.3 | 14.1 | 19 | 21 | 22.9 | 23.3 | 24.2 | 25.3 |
| **HEPES** | 20 | 20 | 20 | 20 | 20 | 20 | 20 | 20 | 20 | 20 | 20 | 20 | 20 |
| **D-Mannitol** | 127 | 120 | 122 | 118 | 116 | 121 | 113 | 108 | 105 | 103 | 103 | 104 | 105 |
| **EGTA** | 25 | - | 25 | 25 | 25 | - | - | - | - | - | - | - | - |
| **HEDTA** | - | - | - | - | - | 25 | 25 | 25 | 25 | 25 | 25 | 25 | 25 |
| **EDTA** | - | 25 | - | - | - | - | - | - |  |  |  |  |  |
| Total [Cl^-^]_i_ | 51.2 | | | | | | | | | | | | |
